# A blood-brain-barrier penetrant AAV gene therapy rescues neurological deficits in mucolipidosis IV mice

**DOI:** 10.1101/2023.11.03.565568

**Authors:** Madison Sangster, Martha Bishop, Yizheng Yao, Jessica Feitor, Sanjid Shahriar, Maxwell Miller, Anil K. Chekuri, Bogdan Budnik, Fengfeng Bei, Yulia Grishchuk

**Affiliations:** Center for Genomic Medicine and Department of Neurology, Massachusetts General Hospital Research Institute and Harvard Medical School, Boston, MA, USA; Department of Neurosurgery, Brigham and Women’s Hospital, Harvard Medical School, Boston, MA; Wyss Institute for Biologically Inspired Engineering, Harvard University, Boston, MA, USA; Grousbeck Gene Therapy Center, Schepens Eye Research Institute, Massachusetts Eye and Ear Infirmary, Harvard Medical School, Boston, MA

## Abstract

Mucolipidosis IV (MLIV) is a rare, autosomal recessive, lysosomal disease characterized by intellectual disability, motor deficits and progressive vision loss. Using AAV9 and AAV-PHP.B as delivery vectors, we previously demonstrated the feasibility of modifying disease course in a mouse model of MLIV by the human *MCOLN1* gene transfer. Here, using a primate-enabling capsid AAV.CPP.16 (CPP16), we constructed a new, clinic-oriented *MCOLN1* gene expression vector and demonstrated its efficacy in the preclinical model of MLIV. Systemic administration of CPP16-*MCOLN1* in adult symptomatic *Mcoln1^-/-^* mice at a dose of 1e12 vg per mouse resulted in *MCOLN1* expression in the brain and peripheral tissues, alleviated brain pathology, rescued neuromotor function, and completely prevented paralysis. Notable expression of *MCOLN1* transcripts was also detected in the retina of the mouse that had exhibited significant degeneration at the time of the treatment. However, no increase of retinal thickness was observed after the gene therapy treatment. Our results suggest a new AAV-based systemic gene replacement therapy for the treatment of MLIV that could be translated into clinical studies.

## Introduction

Mucolipidosis IV (MLIV) is a rare pediatric neurologic disease caused by loss-of-function mutations in the *MCOLN1* gene (1). MLIV was first described in 1974 (2), and the causative gene was identified in 1999 (3, 4, 5). Patients typically present with corneal clouding and delayed developmental milestones in the first year of life and reach a plateau in psychomotor development by two years of age. Early onset of axial hypotonia and signs of pyramidal and extrapyramidal motor dysfunction prevent independent ambulation in the majority of MLIV patients and severely limit fine motor function. Though MLIV was originally described as a static neurodevelopmental disorder, progressive neurological deterioration has recently been documented during the second decade of life (6). In congruence with the clinical course, brain imaging has demonstrated stable white matter abnormalities (corpus callosum hypoplasia and dysgenesis, and white matter lesions) with the emergence of subcortical volume loss and cerebellar atrophy in older patients (7, 8). Visual impairment is also a prominent feature of MLIV with progressive retinal dystrophy and optic nerve atrophy leading to blindness by the second decade of life (8, 9, 10, 11), further impeding function and negatively impacting quality of life. At present, the standard of care of MLIV primarily focuses on symptom management, and no disease modifying treatments are available.

*MCOLN1* encodes the late endosomal/lysosomal non-selective cation channel TRPML1, that regulates lysosomal ion balance and is directly involved in multiple lysosome-related pathways, including Ca^2+^-mediated fusion/fission with the lysosomal membrane, mTOR-signaling, TFEB activation, lysosomal biogenesis (12, 13, 14, 15), and autophagosome formation (16). Additionally, its role in Fe^2+^-transport and regulating brain iron homeostasis has also been demonstrated (17, 18).

Important insights into the pathophysiology of the disease have been obtained using the *Mcoln1* knock-out mouse model we developed (19, 20, 21, 22). *Mcoln1^-/-^* mice recapitulate the clinical and pathological phenotype of MLIV patients including motor deficits, retinal degeneration, corpus callosum hypoplasia, microgliosis, astrocytosis and, later in disease, partial loss of Purkinje cells.

Recently we showed that systemic *MCOLN1* gene transfer using AAV-PHP.B, a mouse-restricted blood-brain barrier (BBB) penetrant capsid (23), can either prevent or fully reverse neurological dysfunction in the MLIV mouse model, demonstrating the feasibility of altering the disease course of MLIV using gene therapy (24). Additionally, intracerebroventricular injection of a self-complementary, *MCOLN1*-expressing AAV9 vector (scAAV9-MCOLN1) in neonatal *Mcoln1^-/-^*pups was also efficacious as wide brain transduction was achieved using this route of administration in young animals. However, when scAAV9-MCOLN1 was intravenously administered in adult symptomatic mice leading to successful overexpression of the transgene in the periphery but little transduction in the brain, no therapeutic effect was observed (24). These studies suggest that broad brain targeting by a human-translatable vector is essential in achieving optimal therapeutic outcomes when developing AAV gene therapy for MLIV.

In this study, we tested the preclinical efficacy of human *MCOLN1* gene transfer using a recently reported BBB-penetrant AAV capsid, AAV.CPP.16 (CPP16). Compared to its parent capsid AAV9, systemic administration of CPP16 demonstrates over 5-fold enhancement in transduction of the CNS in both mice and non-human primates (25). Although it may not be the most potent in overcoming the mouse BBB as compared with other recently developed capsids (26, 27), the translatability of CPP16 from rodents to primates prompted us to examine whether systemic delivery of CPP16-*MCOLN1* would restore sufficient *MCOLN1* expression in the brain and yield meaningful functional outcome. Indeed, we found that intravenous delivery of this vector in symptomatic *Mcoln1^-/-^* mice led to dose-dependent improvement of motor function, significantly delayed time to paralysis, and corrected brain pathology in treated *Mcoln1^-/-^* animals. These data suggest that AAV.CPP.16-mediated systemic gene replacement therapy could be a promising approach for treating patients with MLIV.

## Results

### Systemic administration of CPP16-*MCOLN1* in young adult symptomatic *Mcoln1^-/-^* mice resulted in dose-dependent restoration of neuromotor function and delayed onset of paralysis

Self-complimentary CPP16 vector for this study was produced using the same *MCOLN1* expression construct that we previously created to package scAAV9-*MCOLN1* vector which showed efficacy in *Mcoln1^-/-^* mice when delivered via neonatal ICV administration (24). To test efficacy of scAAV-CPP16-MCOLN1, cohorts of male and female *Mcoln1^-/-^* and wild-type littermate control mice received tail-vein injections of either 5e11 vg, 1e12 vg of CPP16-MCOLN1 or saline at the age of 2 months, when the *Mcoln1^-/-^* mice develop decline of motor function in the form of vertical activity. Efficacy was assessed using our established standard outcome measures including open field and rotarod tests, assessment of clasping and righting reflex. The experimental design including group size and order of testing is shown in **Figure 1**. Reduction of vertical activity is one of the first signs of motor function decline in *Mcoln1^-/-^* mice that can be measured in the open field test starting at 2 months of age (24, 28). Consistent with our previous findings, significant reduction of vertical movements and time spent in the vertical position was observed in saline-treated *Mcoln1^-/-^*mice compared to their WT littermates in both sexes at the age of 4 months **(Figure 2A and B)**. In females, intravenous administration of CPP16-MCOLN1 at 1e12 vg/animal led to significant rescue of vertical movements and vertical time indicating the restoration of neurologic function **(Figure 2A)**. In males, we observed a similar trend towards higher vertical activity in *Mcoln1^-/-^* animals treated with either 5e11 or 1e12 vg/mouse of CPP16-MCOLN1 as compared to saline-treated *Mcoln1^-/-^*group. Statistical significance between saline and CPP16-MCOLN1-treated *Mcoln1^-/-^* males was not detected likely due to small sample size.

**Figure 1.**
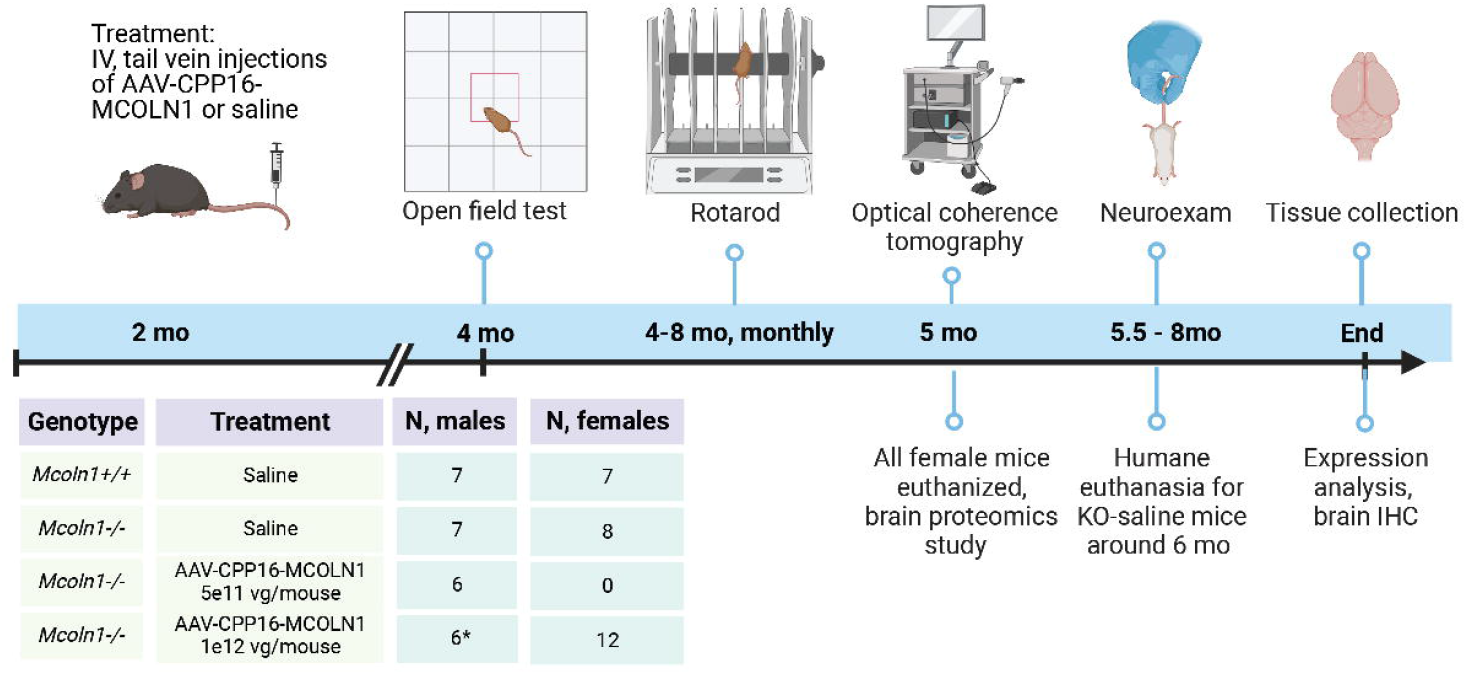
Schematic representation of CPP16-MCOLN1 efficacy study in *Mcoln1^-/-^* mice and study groups. * - one male developed severe health concern (penile prolapse) and had to be euthanized at 3 months of age. Assessment of efficacy after 3 months of age is reported for n=5 mice in WT saline group.

**Figure 2.**
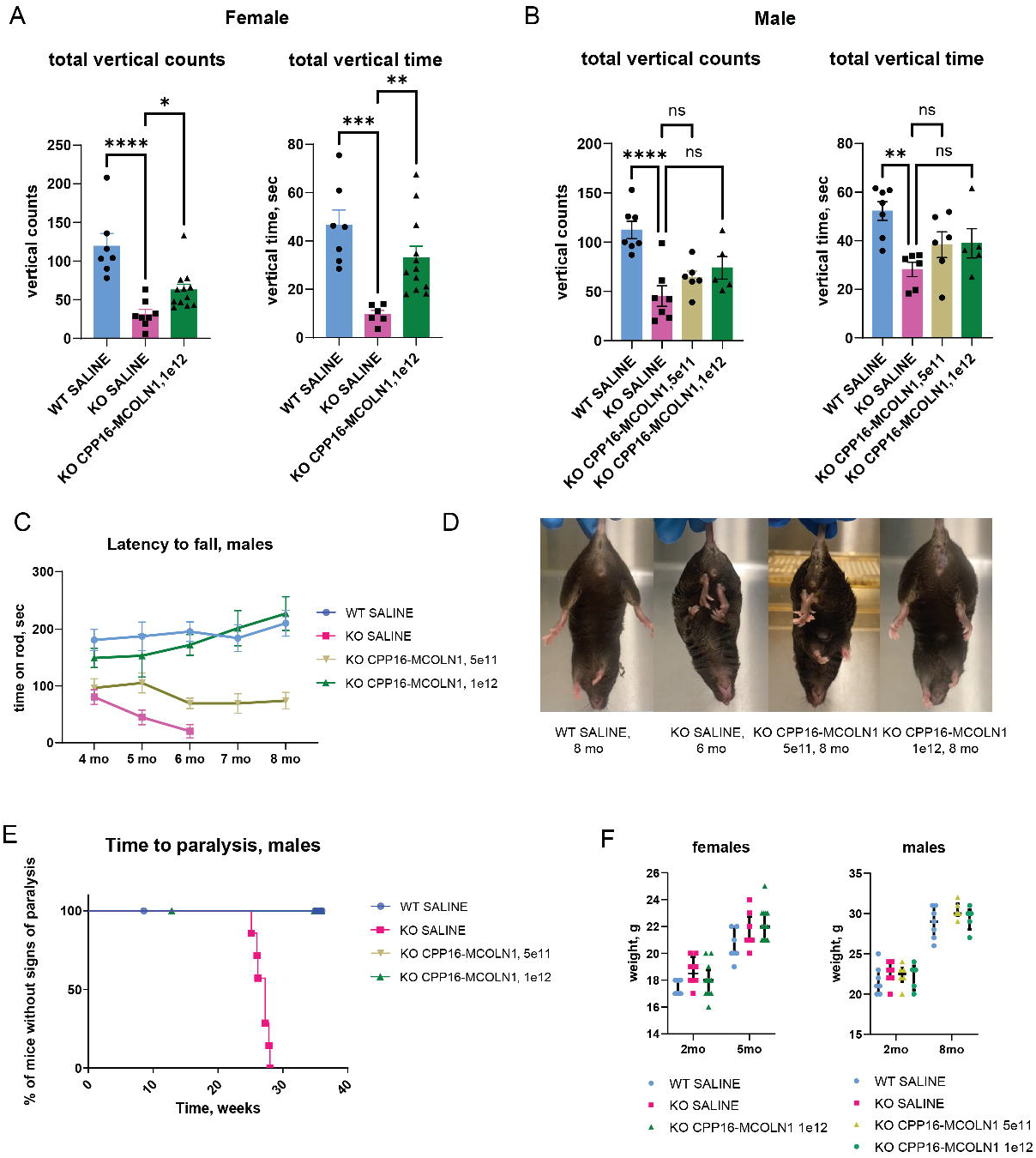
Delayed paralysis and dose-dependent restoration of neurologic function after intravenous administration of CPP16*-MCOLN1* in symptomatic *Mcoln1^-/-^* mice. **A, B.** Measurements of the vertical activity in the open field test, represented as total vertical counts and total vertical time in the open field arena, in female (A) and male (B) mice show significantly decreased activity in the 4 months-old *Mcoln1^-/-^* mice treated with saline compared to saline-treated WT controls, and significant recovery in the *Mcoln1^-/-^* mice that were treated with CPP16-*MCOLN1* at the symptomatic stage of the disease at 2 months of age. Data presented as individual data points per animal, mean values, and SEM; two female and one male mice were identified as outliers and excluded from stat analysis; group comparisons made using one-way ANOVA test; represented p values were corrected for multiple comparisons between individual groups. **C.** Dose-dependent long-term improvement of the rotarod performance presented as average latency to fall indicates better motor function, balance and coordination in the *Mcoln1^-/-^* mice treated with CPP16-*MCOLN1.* **D.** Representative images of mice in all treatment/genotype groups showing rescue of clasping in the *Mcoln1^-/-^*mice treated with 1e12 vg/mouse, but not with 5e11 vg/mouse of CPP16-*MCOLN1.* **E.** Systemic administration of CPP16-*MCOLN1* at 2 months of age significantly delays time to paralysis in *Mcoln1^-/-^* male mice. The criterion for paralysis was loss of righting reflex when mouse failed to rotate itself in upright position after placing on a side within 10 sec; log-rank test p-value is <0.0001. **F.** No significant weight changes have been observed in mice treated with CPP16-*MCOLN1*. Data presented as median values and interquartile range.

Only the male cohort of mice were used for the rotarod testing. Compared to healthy littermates, saline-treated *Mcoln1^-/-^* mice showed a lower average latency to fall starting at the age of 4 months **(Figure 2C).** Gradually, saline-treated *Mcoln1^-/-^* mice developed hind limb weakness and were eventually euthanized at around 6 months of age due to hind-limb paralysis. Importantly, *Mcoln1^-/-^* mice that received 1e12 vg of CPP16-MCOLN1 (high dose) showed comparable performance on rotarod with the control healthy littermates up to the end of the trial at 8 months of age. Furthermore, no hind limb clasping was observed in these animals and their overall appearance was undistinguishable from the saline-treated wild-type mice **(Figure 2D)**. *Mcoln1^-/-^* group treated with 5e11vg of CPP16-*MCOLN1* per animal (low dose) did not perform as well on rotarod as the animals in the high-dose group, showing lower rod-retention time **(Figure 2C)**. The low-dose group also showed no rescue of clasping **(Figure 2D)** and had scruffy coat appearance resembling saline-treated *Mcoln1^-/-^*mice. Interestingly, despite the general ill appearance of the *Mcoln1* knock-out mice treated with the low dose of AAV-CPP16-*MCOLN1,* none of these mice developed signs of hind-limbs paralysis throughout the 8-month study, showing extended lifespan by at least 2 months over saline-treated *Mcoln1^-/-^*mice **(Figure 2E).** We observed no weight changes or health concerns in the CPP16-MCOLN1 treated mice **(Figure 2F).** In conclusion, our data show a dose-dependent response of CPP16-MCOLN1 gene therapy in MLIV with neuromotor functions fully rescued by a single, high-dose, systemic administration.

### Expression of *MCOLN1* transgene in the CNS and peripheral tissues

q-RT-PCR analysis of the *MCOLN1* transgene expression in post-mortem tissues showed a dose-dependent increase of mRNA transcripts in the brain regions (cortex and cerebellum), retina, sciatic nerve, skeletal muscle, and stomach of *Mcoln1^-/-^*males treated with 5e11 or 1e12 vg/mouse of CPP16-*MCOLN1* **(Figure 3A).** No significant differences were detected between male and female groups treated with CPP16-*MCOLN1* at 1e12 vg/mouse, except in the retina, where higher expression was detected in female mice. Notably, transduction and transgene expression in the retina with systemic administration of CPP16 capsid has not been reported previously. In line with the previously published data of CPP16 biodistribution in the mouse tissues (25), we observed high expression of *MCOLN1* transcripts in the liver. Remarkably high expression was also detected in the skeletal muscle.

**Figure 3.**
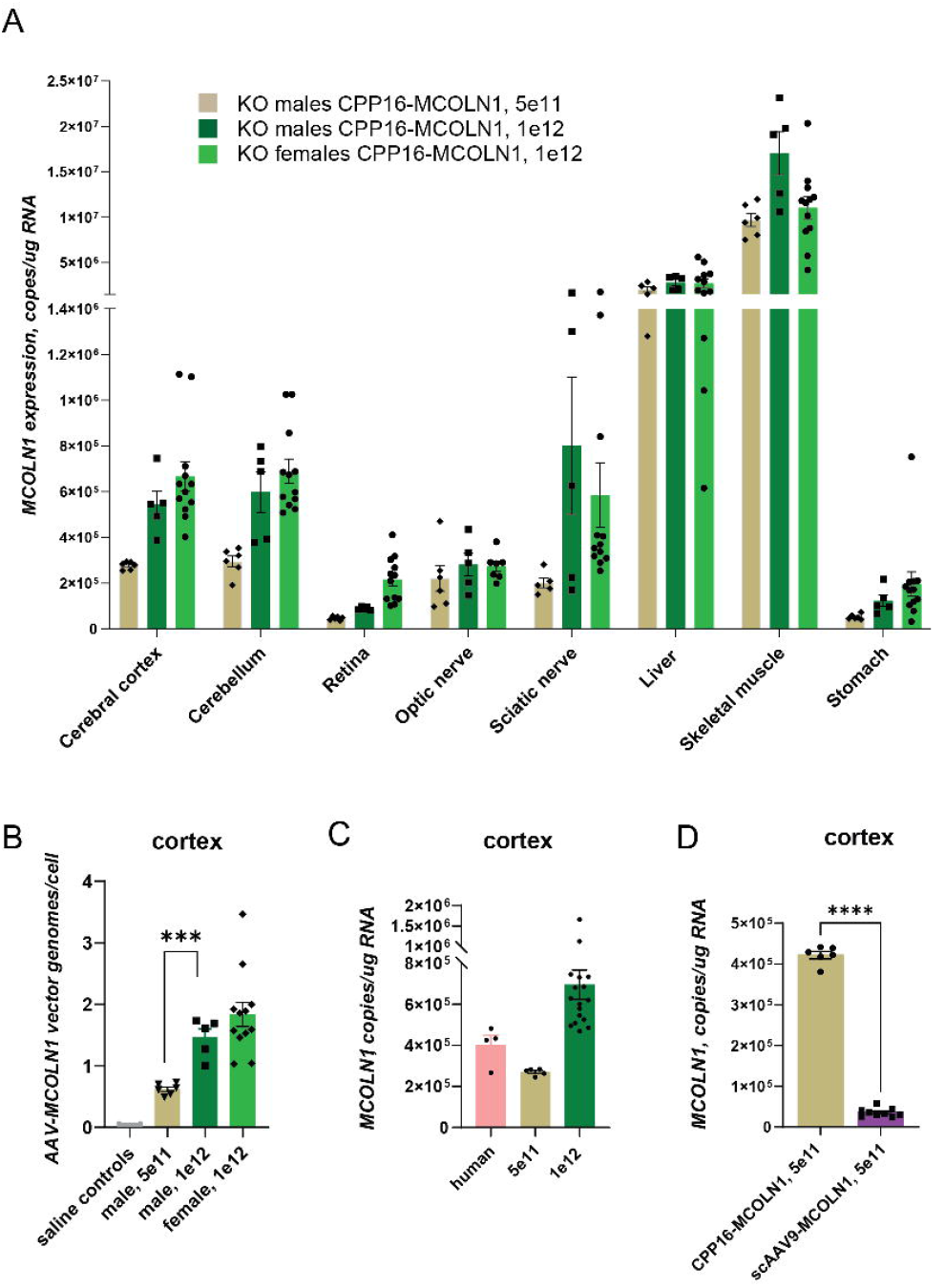
*MCOLN1* expression analysis in post-mortem mouse tissues shows transduction in CNS and peripheral tissues. **A.** qRT-PCR analysis of the *MCOLN1* transcripts showing transgene overexpression in the cerebral cortex, cerebellum, retina, optic nerve, sciatic nerve, liver, quad muscle, and stomach. **B.** qPCR vector genome copy analysis showing CPP16-*MCOLN1* vector biodistribution in the cerebral cortex after intravenous administration to 2 months-old mice. **C.** qRT-PCR analysis showing AAV-CPP16-driven human *MCOLN1* expression in the mouse cortex vs. endogenous *MCOLN1* expression in the human cortex. P (one-way ANOVA) =0.0048. **D.** qRT-PCR analysis in *Mcoln1^-/-^* mice injected intravenously with 5e11vg/animal of either self-complimentary CPP16-*MCOLN1* or AAV9-*MCOLN1* shows that CPP16 is superior in transducing the brain from systemic flow as compared to scAAV9. Data presented as individual data point, mean values, and SEM. P, unpaired T-test, *** is <0.001; **** is <0.0001.

Vector genomes quantification analysis in the cerebral cortex showed a dose-dependent increase of viral genome copies per cell in the male groups from the mean of 0.61 (± 0.037, SEM) in the 5e11 vg/mouse group to 1.46 (± 0.14) in the 1e12 vg/mouse group **(Figure 3B)**. An average of 1.83 (± 0.195) vg/cell was detected in the female mice treated with 1e12 vg of the vector. This data, together with the motor function outcomes described earlier, suggests that average cortical biodistribution higher than 0.61 vg/cell is required to obtain restoration of the neurologic function in MLIV.

To assess how therapeutic levels of CPP16-mediated *MCOLN1* expression determined in the mouse tissue are compared to amount of endogenous human *MCOLN1* expression, we measured *MCOLN1* mRNA in 4 human cortical samples (young adult male; Maryland Brain Biobank) and compared them with the human MCOLN1 transgene mRNA levels in the cortices of the *Mcoln1^-/-^* mice treated with AAV-CPP16-MCOLN1 **(Figure 3C)**. We found that treatment with the low dose of CPP16-*MCOLN1* (5e11 vg per mouse) resulted in expression of *MCOLN1* mRNA lower than the endogenous level in the human cortex, while treatment of the high dose (1e12 vg per mouse) led to, on average, higher than endogenous level *MCOLN1* transcript expression.

The superiority of CPP16’s ability to transduce brain tissue from systemic flow over AAV9 has previously been demonstrated (25). To confirm this in our study, we performed direct comparison of *MCOLN1* mRNA levels in the cerebral cortex tissue of *Mcoln1^-/-^* mice that received either self-complimentary AAV9 or self-complimentary CPP16 vectors carrying the same expression cassette via the same route of administration (tail-vein injection). Consistent with the previous report, we found CPP16-mediated *MCOLN1* expression was 12-fold higher than that achieved by AAV9 **(Figure 3D).**

### Systemic treatment with CPP16-*MCOLN1* did not alleviate retinal pathology

Eye pathology in *Mcoln1^-/-^* mice includes thinning of the photoreceptor layer, reduced levels of rhodopsin, and significantly decreased dark-adapted a- and b- response indicative of the rod cells impairment (21). Remarkably, this retinal phenotype in *Mcoln1^-/-^* mice was found to be present early, at 1 month of age, and was static, in contrast to progressive retinal, corneal and optic nerve pathology in MLIV patients that leads to complete blindness in the second decade of life. Since our expression data showed expression of the human *MCOLN1* transgene in the *Mcoln1^-/-^* mouse retina after intravenous administration of CPP16-MCOLN1, we set out to assess whether this expression resulted in correction of retinal thinning. Retinal thickness was measured in live mice using high-definition spectral domain optical coherence tomography (SD-OCT). Male 5-month-old WT and *Mcoln1^-/-^*mice that received either saline or 1e12 vg of CPP16-*MCOLN1* were used. The SD-OCT showed significant retinal thinning, with a reduced thickness of the photoreceptor (outer nuclear) layer and the outer photoreceptor segments in saline-treated *Mcoln1^-/-^* mice compared to saline-treated WT, confirming retinal degeneration. Unfortunately, no improvement of the retinal or outer nuclear layers was detected in the *Mcoln1^-/-^* mice treated with CPP16-*MCOLN1* **(Supplementary Figure 1)**. Given that retinal phenotype is fully developed in the *Mcoln1^-/-^*mice by 1 month of age and CPP16-*MCOLN1* in our study was administered when mice reached 2 months of age, the lack of therapeutic benefit may be a result of belated intervention.

### CPP16-*MCOLN1* reduces brain pathology in *Mcoln1^-/-^* mice

An increased abundance of the lysosomal proteins and decreased levels of oligodendrocyte and myelin related proteins are molecular hallmarks in the brain of symptomatic *Mcoln1^-/-^*mice (29). To assess if administration of CPP16*-MCOLN1* rescued brain pathology in *Mcoln1^-/-^* mice, we next performed LC-MS/MS proteomics analysis using cortical tissues of 5-month-old female *Mcoln1^-/-^*mice as well as wild-type control mice. In accordance with previous findings, we observed a remarkably similar broad upregulation of the lysosomal proteins and downregulation of myelination and oligodendrocyte protein signature in the *Mcoln1^-/-^*cortex **(Figure 4A).** In total, 3490 proteins were identified **(Table S1)**. Mouse albumin and keratin entries and non-mouse proteins were removed from these data sets (the list of all typical contaminants is available in **Table S2**). Principal component analysis (PCA) separated *Mcoln1^-/-^* - saline, *Mcoln1^-/-^* -CPP16-MCOLN1 and WT-saline samples **(Figure S2)**. 28 upregulated and 31 downregulated proteins were detected in the cortical tissue of saline-treated *Mcoln1^-/-^* mice as compared to WT-saline group, with log2 foldchange cutoff at 0.5 (fold change of +/-1.4) and log10 p-value cut off at 1.3 (p-value <0.05) **(Figures 4A, S3 and Table S3)**. 18 out of 28 upregulated proteins were lysosomal. Remarkably, we observed a broad reduction of lysosomal protein levels in the *Mcoln1^-/-^* mice treated with CPP16-MCOLN1. Selected examples including Protein phosphatase 1 regulatory subunit 21 (Ppp1r21), Beta-hexosaminidase subunit B (Hexb), Arylsulfatase B (Arsb), Cathepsin D (Ctsd), and Ganglioside GM2 activator (Gm2a) are presented in **Figure 4B**. In addition, we observed a reduction of the astrocytosis marker Glial fibrillary acidic protein (Gfap) and inflammation-linked proteins including Platelet-activating factor acetylhydrolase (Pla2g7), Signal transducer and activator of transcription 1 (Stat1), 5’-3’ exonuclease PLD3 (Pld3) in CPP16-*MCOLN1*-treated samples. This data demonstrates at least partial correction of lysosomal and pro-inflammatory phenotype in the *Mcoln1^-/-^* mouse brain following gene transfer of human *MCOLN1*. Our LC-MS/MS data showed no recovery of myelin and oligodendrocyte related proteins in *Mcoln1^-/-^* mice treated with CPP16-*MCOLN1*. The complete proteomic data set is presented in **Table S4**.

**Figure 4.**
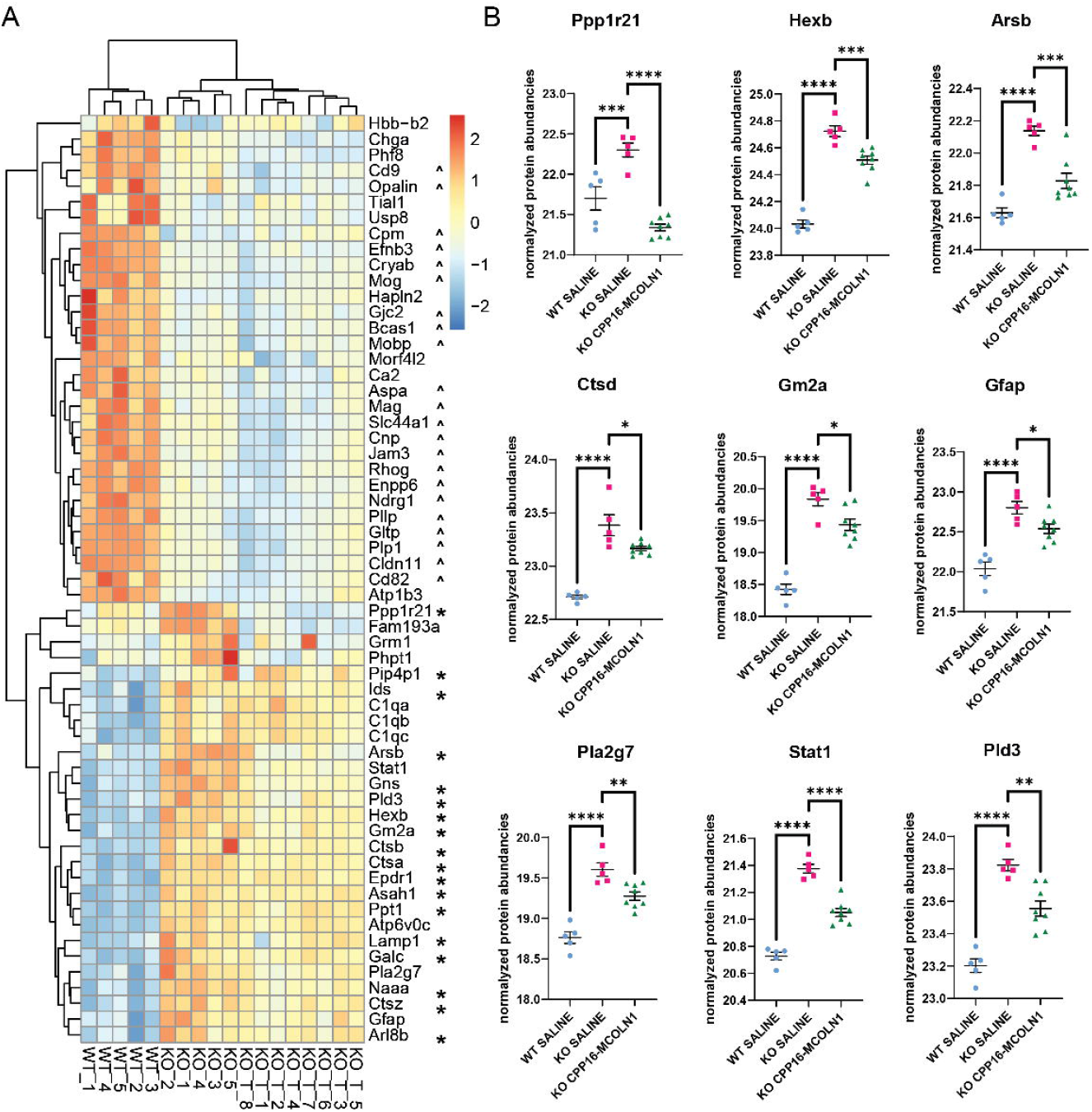
Intravenous administration of CPP16-*MCOLN1* in symptomatic *Mcoln1^-/-^* mice partially corrects upregulation of lysosomal proteins in the brain. **A.** Heatmap showing downregulated and upregulated proteins (log10 p<0.05, log2FC>0.5) in *Mcoln1^-/-^*-saline cortical homogenates from 5-month-old female mice compared to WT-saline littermates and corresponding protein abundancies in *Mcoln1^-/-^* CPP16-*MCOLN1*-treated mice. * mark lysosomal proteins; ^ mark proteins enriched in oligodendrocyte cell lineage and myelin. **B.** Individual proteins in endosomal/lysosomal compartment (upper panel) or glial and immune-related proteins, demonstrating significant increase in saline-treated KO and their correction in KO mice treated with CPP16-*MCOLN1*. P, one-way ANOVA and Dunnett test for multiple comparisons, *<0.05, **<0.01, ***<0.001, ****<0.0001. See also Figures S2 and S3 and tables S1-S4.

To confirm our observation of lysosomal protein correction using LC-MS/MS proteomics, we performed immunohistochemistry to examine lysosomal pathology. Lamp1 is broadly used as lysosomal marker and increases in percentage of Lamp1+ staining and size of Lamp1+ particles are indicative of lysosomal abnormality in *Mcoln1^-/-^*mice (24). We found that intravenous administration of AAV-CPP16-MCOLN1 vector led to significant reduction of both % of stained area and particle size measurements (**Figure 5A)**, demonstrating reduction of the lysosomal pathology in the treated *Mcoln1^-/-^* group.

**Figure 5.**
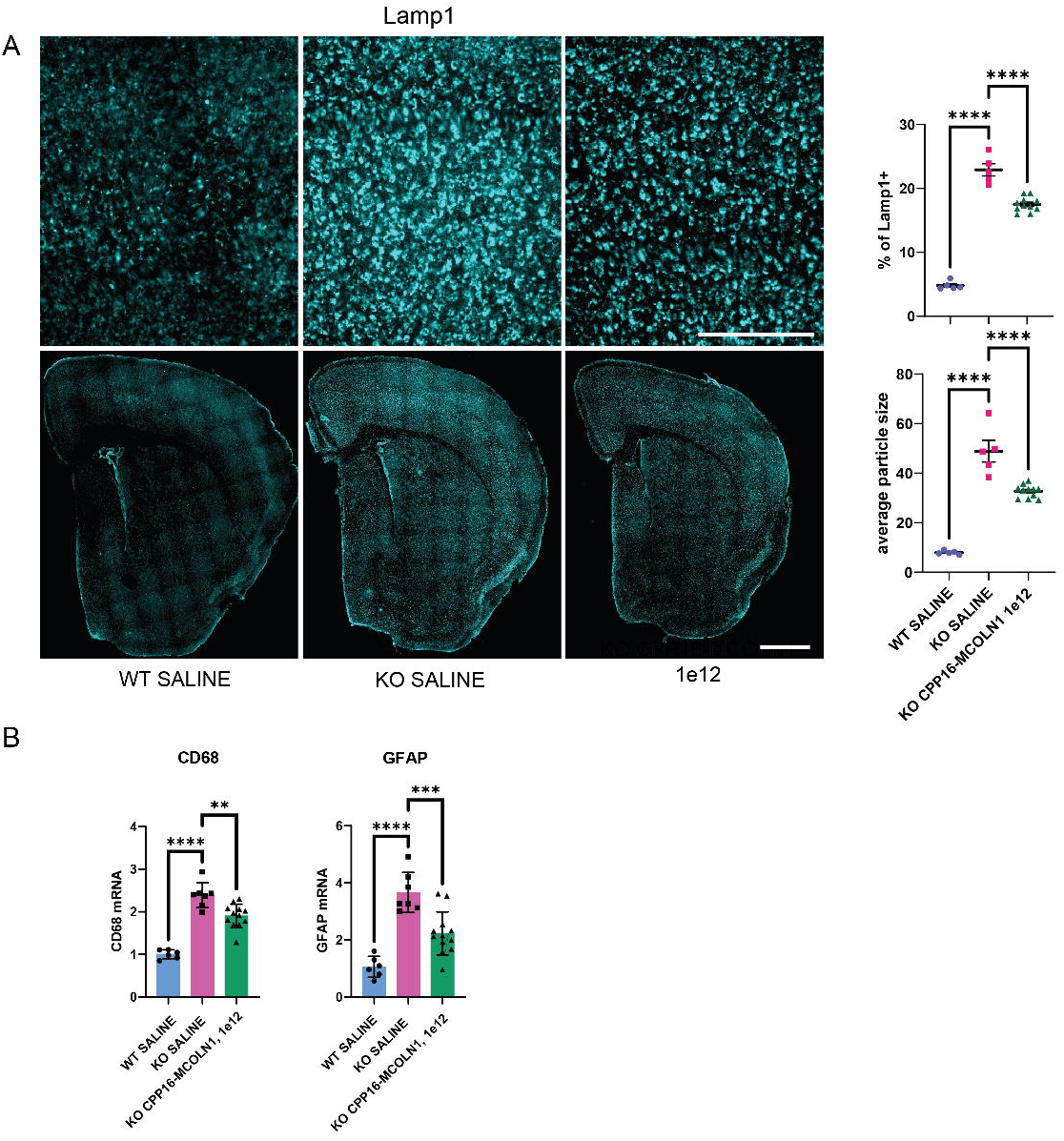
Intravenous administration of CPP16-*MCOLN1* in symptomatic *Mcoln1^-/-^* mice reduces accumulation of Lamp1-positive lysosomal aggregates and reduces activation of astrocytosis and microgliosis in *Mcoln1^-/-^*mouse brain. **A.** Representative images of Lamp1 staining in mouse brain at higher (upper panel, scale =0.25 mm) and lower (lower panel, scale = 1mm) magnification, and image quantification showing average Lamp1^+^ particle size and % of Lamp1^+^ staining area in the coronal whole hemisphere sections. Data show individual, mean values and SEM, n (WT-saline, female) = 5; n (*Mcoln1^-/-^* saline, female) = 5; n (*Mcoln1^-/-^*AAV-CPP16-MCOLN1, 1e12, female) = 11. Comparisons done using ordinary one-way ANOVA with Dunnett’s test for multiple comparisons using GraphPad Prism v.9; p, **<0.01, ****<0.0001. **B.** qRT-PCR analysis of the microglia activation marker CD68 and astrocytosis marker GFAP showing significant upregulation of the corresponding transcripts in the cerebral cortex of *Mcoln1^-/-^* mice treated with saline compared to saline-treated WT littermate mice and significant reduction of the CD68 and GFAP transcripts in the *Mcoln1^-/-^* mice treated with CPP16-*MCOLN1*. Data show individual, mean values and SEM, n (WT-saline, female) = 6; n (*Mcoln1^-/-^*saline, female) = 7; n (*Mcoln1^-/-^* CPP16-*MCOLN1*, 1e12, female) = 11. Comparisons done using ordinary one-way ANOVA with Dunnett’s test for multiple comparisons using GraphPad Prism v.9; p, **<0.01, ***<0.001 ****<0.0001.

We further performed qRT-PCR analysis to examine astrocytosis and microgliosis that are early hallmarks of the MLIV brain pathology reported in the human tissue and the MLIV mouse model (28, 30, 31, 32). As expected, significant increases of mRNA transcripts of the astrocytosis marker *Gfap* and microgliosis marker *Cd68* were observed in the cortical tissue of saline-treated *Mcoln1^-/-^* mice compared to saline-treated WT littermates **(Figure 5B)**. Levels of both transcripts were significantly reduced in the *Mcoln1^-/-^* mice treated with CPP16-*MCOLN1* indicating reduction of glial pathology **(Figure 5B)**. An observed reduction of GFAP transcripts in *Mcoln1^-/-^* mice after treatment with CPP16-*MCOLN1* is consistent with our observation on reduced GFAP protein abundance using LC-MS/MS **(Figure 4B)**.

### Pierce correlation analysis of efficacy outcomes and brain expression of *MCOLN1* transgene

Pierce correlation analysis in the male and female cohorts showed that higher *MCOLN1* transgene expression in the cerebral cortex correlated with higher expression in the cerebellum as well with better performance in the rotarod test and lower expression of the glial pathology markers, GFAP (males) and CD68 (females) **(Figure 6)**, supporting the therapeutic effect of *MCOLN1* gene replacement in the *Mcoln1^-/-^* mice.

**Figure 6.**
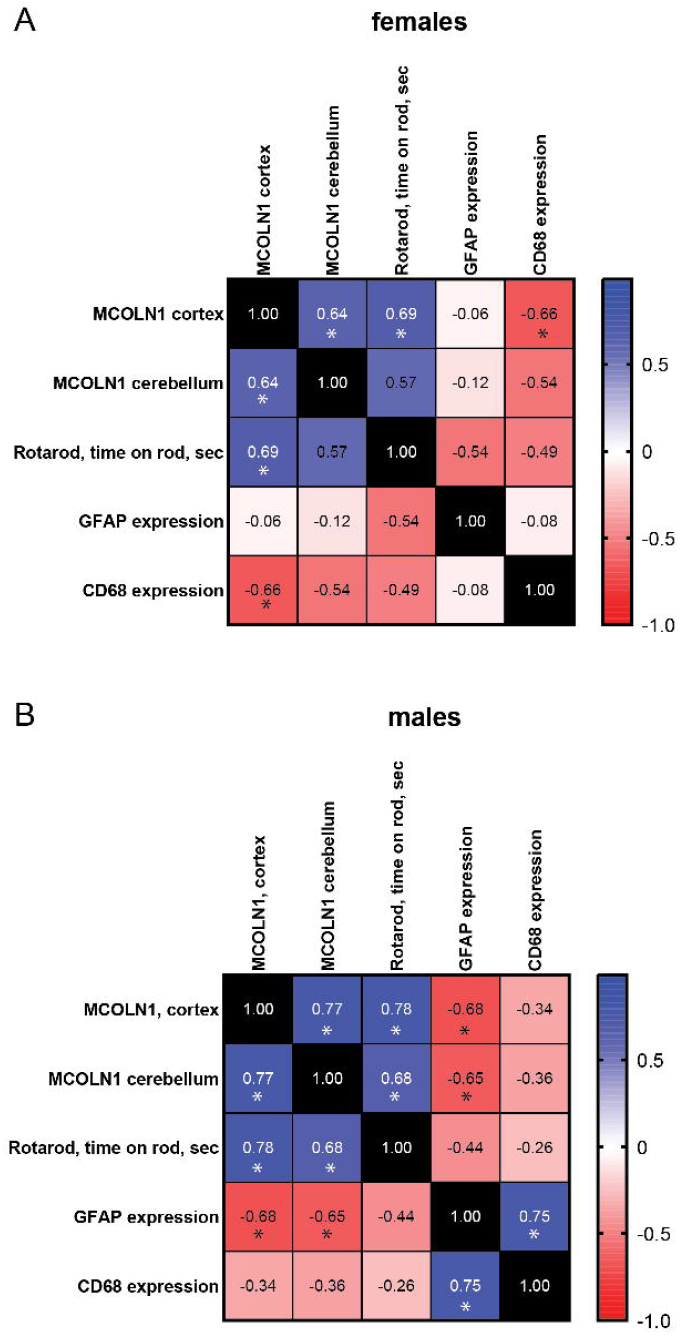
Pierce correlation analysis of efficacy outcomes and brain expression of *MCOLN1* transgene. Heatmaps show Peirce correlation coefficient between *MCOLN1* transgene expression in either cerebral cortex or cerebellum and efficacy outcomes, including rotarod performance and expression of glial markers, GFAP and CD68. Asterisks show p-values <0.05. Data in female cohort is shown in **A**, data in male cohort is shown in **B**.

## Discussion

Gene delivery to sufficient percentages of cells across broad regions in the CNS has been a major hurdle preventing development of a gene therapy for MLIV, a disease with high unmet need and critical CNS manifestations. Systemic route of administration via the bloodstream would be an ideal approach for delivering a therapy to both CNS and peripheral targets. While AAV9 is the AAV serotype of broad tissue tropism and a proven track record in human application, its CNS delivery efficiency is still suboptimal because of the limitation posed by the BBB. Extensive efforts have been made in engineering a new generation of AAV variants with different extent of BBB penetrance such as AAV-AS, Anc80L65, AAV-PHP.B, AAV-F, 9P31 and AAV.CAP-Mac (26, 33, 34, 35, 36). However, it is increasingly evident that genetic drift between species poses a challenge when using AAVs selected based on small animal models for human applications. For example, study with AAV-PHP.B, which is over 40-fold more efficient than AAV9 in overcoming the BBB in C57BL/6 mice, does not translate into primates because of differential expression of its mouse-restricted receptor LY6A (37, 38). We previously used AAV-PHP.B to systemically deliver a supra-physiological level of the *MCOLN1* transcript to the brain in *Mcoln1^-/-^* mice and observed correction of neurological dysfunction (24). Lack of translatability for AAV-PHP.B prompted us to test CPP16, the superiority of which over AAV9 translates from mice to NHPs (25). CPP16 was developed through a rational-design approach by inserting a cell-penetrating-peptide (CPP) derived peptide (“TVSALK”) to the aa588/589 site of AAV9 capsid. Although the molecular mechanism underlying the enhanced neurotropism of CPP16 over AAV9 remains to be unveiled, increased transcytosis in the brain microvascular endothelial cells in a human BBB model and better transduction of human brain cells have been observed (25). Thus, CPP16 has the potential for translation into clinical testing. The fact that systemic application of CPP16-mediated *MCOLN1* gene transfer was efficacious in the mouse model, as we showed in this study, provides additional evidence supporting further development of such therapy for MLIV patients.

Biodistribution of CPP16 and transgene expression outside of the CNS were not systemically investigated in the previous study (25). Our *MCOLN1* expression data after intravenous administration of CPP16-*MCOLN1* in young adult mice showed broad biodistribution and expression of the transgene not only in the brain tissue, but also in the retina, optic nerve and other peripheral nerve tissues, as well as in other peripheral organs such as the skeletal muscle and stomach. Given the complex clinical presentation of MLIV, including retinal degeneration, optic nerve pathology, malfunctioning of parietal cells in the stomach leading to achlorhydria, therapeutic approach involving systemic administration of CPP16-*MCOLN1* may simultaneously target the CNS, the eye and peripheral organs, eliciting maximal benefits in MLIV in the clinical setting. Since many genetic syndromes with early onset and CNS involvement are characterized by peripheral pathology and visual abnormalities, including many lysosomal, mitochondrial diseases and metabolic syndromes, CPP16 could present an attractive vector platform with applications beyond MLIV.

Our study was designed to assess efficacy of the *MCOLN1* gene transfer in early symptomatic *Mcoln1^-/-^*mice. The timing of intervention was selected to closely match the design for future clinical studies. Due to early onset of the disease in humans, a vast majority of patients with MLIV have developed neurological symptoms by the time they are diagnosed. MLIV families and caregivers identify early motor dysfunction as a major contributor to disability and limited quality of life in MLIV patients. Thus, a successful treatment for MLIV ought to rescue developmental motor deficits rather than delay motor deterioration. Motor deficits in *Mcoln1^-/-^* mice first appear at the age of two months in the form of reduced rearing or vertical activity, and then progress gradually to loss of ambulation due to hind-limb paralysis and premature death by 7 months of age (20), (24). Our observation of therapeutic benefits in early symptomatic mice after treatment with CPP16-based gene therapy suggests MLIV patients may also benefit from this approach.

Dose determination is an essential step in designing future clinical trials. We observed that both doses, 5e11 and 1e12 vg/mouse, were able to delay time of paralysis and extend lifespan of the treated *Mcoln1^-/-^*mice and that only the higher dose of 1e12 vg/mouse (equivalent to ∼5e13 vg/kg) led to significant restoration of the neurologic function, improvement of the rotarod performance, full rescue of the clasping phenotype, and the overall healthy appearance. The observed dose dependence of CPP16-*MCOLN1* therapy provides important guidance in selecting a range of doses for clinical testing. We further observed that a full therapeutic dose, 1e12 vg/mouse, of CPP16-*MCOLN1* corresponds to vector biodistribution of approximately 1.7 vg/cell in the cortex. Since vector biodistribution of AAVs can be measured relatively easily across species, determination of the biodistribution threshold/target for CPP16-MCOLN1 in our study provides a biomarker for follow-up translational and clinical studies.

In conclusion, we report a systemic AAV-based gene therapy for MLIV with the promise of translation into clinical setting.

## Materials and Methods

### Animals

*Mcoln1*^-/-^ mice were maintained as previously described (20). Genotyping was performed by Transnetyx using real-time qPCR (transnetyx.com). The *Mcoln1*^+/-^ breeders for this study were obtained by backcrossing onto a C57BL/6J background for more than 10 generations. Experimental cohorts were obtained from either *Mcoln1*^+/-^ x *Mcoln1*^+/-^ or *Mcoln1*^+/-^ x *Mcoln1*^-/-^ mating. *Mcoln1*^+/-^ and *Mcoln1*^+/+^ littermates were used as controls. Experiments were performed according to the Institutional and National Institutes of Health guidelines and approved by the Massachusetts General Hospital Institutional Animal Care and Use Committee. Animals were assigned to the experimental groups in a random order. Handling and testing were performed by investigators blinded to treatment.

### Virus preparation and titration

The previously described expression plasmid pAAVsc-JeT-MCOLN1-pA (24) was used to generate scAAV-CPP16-MCOLN1 used in this study. AAV vector production was done as previously described (25). Briefly, HEK293T cells were co-transfected by three plasmids: a pAAV-RC, a pHelper (240071-12, Agilent Technologies, Santa Clara, CA, USA), and an ITR-flanked AAV plasmid (pAAVsc-JeT-MCOLN1-pA) using polyethyleneimine (PEI) (Cat. no 23966, Polysciences, Warrington PA, USA). AAV was collected from serum-free culture media at 72 h and from the cells and media 120 h post transfection. Viral particles were purified by iodixanol gradient (15%, 25%, 40% and 60%) ultracentrifugation, then concentrated and desalted using 100K Millipore Amicon filter unit (UFC910008, 100 K MWCO), and formulated in Dulbecco’s phosphate buffered saline (PBS). Any viral particle precipitation was re-suspended before application. AAV titers were determined using quantitative PCR. Briefly, AAV samples were treated with DNase I to remove contaminating DNA, followed by sodium hydroxide treatment to lyse the AAV capsid and release the scDNA. pAAVsc-JeT-MCOLN1-pA plasmid was digested with EcoRI (NEB) to recover the ITR-flanked AAV genome to be used as standard. Quantitative PCR was performed with primers targeting the MCOLN1 transgene: forward primer 5′-CAGCACGGAGACAACAGCTT and reverse primer 5′-CAGGGAGCAGGTGAGGATGA.

### IV injections

Mice were restrained in a plexiglass container and the tail veins were dilated under a heat lamp for one minute. 30G-needle insulin syringes (cat. No 328466, BD, San Diego, CA) were used to inject 150-200ul of either 0.9% saline (Hanna’s Pharmaceutical Supply, Wilmington, DE), 5e11 vg scAAV-CPP.16-JeT-*Mcoln1*, or 1e12 vg scAAV-CPP.16-JeT-*Mcoln1* via the tail veins of each mouse.

### Behavioral testing

Open field testing was performed at four months of age under regular light conditions. Testing was done at the same time each day to minimize effects from the circadian cycle, and males were tested the day before females. Each mouse was placed in the center of a 27 × 27 cm^2^ Plexiglass arena, and the horizontal and vertical activities were recorded by the Activity Monitor program (Med Associates Fairfax, VT). Data were analyzed during the first 15 minutes in the arena.

Motor coordination and balance were tested on an accelerating rotarod (Ugo Basile, Gemonio, Italy). Latency to fall from the rotating rod was recorded in 3 trials on one day (accelerating speed from 4 to 40 rpm over 5 min) following a training day of two trials. Animals were tested monthly starting from 4 months of age to 8 months of age or until they reached the euthanasia endpoint such as complete paralysis.

To visualize the clasping reflex in the mice, one investigator held each mouse by the tail while another took a two-second-long video. Screengrabs of the videos are displayed in Figure 2. To determine the euthanasia timepoint for the knockout mice the righting reflex of each mouse was evaluated every two-three days starting at 5.5 months of age. When mice could no longer right themselves in fewer than ten seconds after being placed on their backs, they were euthanized.

### Optical coherence tomography (OCT)

In-vivo imaging of the retina via OCT was performed as described previously (39). Mice were anesthetized in a mobile isoflurane induction chamber with 2% of isoflurane at 2 L/min O2. Pupils of the mice were dilated using 2.5% phenylephrine and 1% tropicamide. About 0.5% of the proparacaine was used as a topical anesthetic. SD-OCT imaging was performed using a Leica EnvisuR2210 (Biooptigen, Durham, United States). Measurements were made 500 μm from the optic nerve for the central retina and 1.5 mm from the optic nerve for the peripheral retina. Linear B-scans of central and peripheral retina were performed, and the thickness of the total retinal, and retinal pigment epithelium (RPE), Outer segments and outer nuclear layer (ONL) layers were measured using Bioptigen Diver software (Bioptigen, Inc., Durham, United States) using automated segmentation. Each OCT image comprises 100 B-scans, with each B-scan consisting of 1000 A-scans. For quantification of retinal thickness, two representative images per eye were analyzed for four images per mouse.

### Tissue collection and processing

Mice were sacrificed using a carbon dioxide chamber. Immediately after euthanasia, mice were transcardially perfused with ice-cold phosphate buffered saline (PBS). The brain was removed and bisected down the midline; one half was post-fixed in 4% paraformaldehyde (Electron Microscopy Sciences, Hatfield, PA) in PBS for 48 hours, washed with PBS, cryoprotected in 30% sucrose in PBS for 24 hours, snap-frozen in isopentane, and stored at −80°C. The other half was further dissected and, along with all other tissues, snap frozen on dry ice.

### RNA extraction and qPCR analysis

Mouse tissues were either homogenized using QIAzol lysis reagent (Qiagen, Hilden, Germany) in a Tissue Lyser instrument (Qiagen, Hilden, Germany), or with a mortar and pestle in buffer RLT (Qiagen, Hilden, Germany) using a 19G needle and syringe. Total RNA isolation from homogenized tissues was performed using Qiagen RNeasy kit (Qiagen, Hilden, Germany) and genomic DNA was eliminated by performing DNase (Qiagen, Hilden, Germany) digestion on columns following the provider’s protocol. cDNA was produced from 500 ng of starting RNA using High-Capacity cDNA Reverse Transcription kit (Applied Biosystems, Foster City, CA). After dilution, 40 ng of the cDNA was used for qPCR using LightCycler 480 Probes Master mix (Roche Diagnostics, Mannheim, Germany), or TaqMan gene expression assays (see details below) on the LightCycler 480 (Roche Diagnostics, Mannheim, Germany). Human tissue was obtained from the NIH NeuroBioBank’s Brain and Tissue repository at the University of Maryland, Baltimore.

To perform relative gene quantification, TaqMan probes (Applied Biosystems, Foster City,CA) were used to measure mouse MBP (FAM)-Mm01266402_m1, mouse GFAP (FAM)-Mm01253033_m1, mouse CD68 (FAM)-Mm03047343_m1, mouse LAMP1 (FAM)-Mm00495262_m1, and human Mucolipin-1 (FAM)- Hs01100653_m1, using mouse GAPDH (FAM)-Mm99999915_g1 as reference. The ΔΔCt method was used to calculate relative gene expression, where Ct corresponds to the cycle threshold. ΔCt values were calculated as the difference between Ct values from the target gene and the housekeeping gene GAPDH.

To perform absolute RNA quantification, three TaqMan probes (APZTMGC) were designed to measure AAV-JeT-driven expression of *MCOLN1* transgene: forward 5’-GGTCGCGGTTCTTGTTTGT-3’, reverse 5’-GAAGCCGCTCGGTCTCT-3’, probe 5’-CCCTGTGATCGTCACTTGACAGTGT-3’. To obtain standard curve, the scAAV-JeT-MCOLN1 plasmid was linearized with HindIII (New England Biolabs, Ipswich, MA, USA) as directed by the manufacturer. The DNA concentration was determined after digestion, and the number of copies of DNA/ul was calculated assuming a molar mass of 650 g/mol per base pair and a fragment length of 5158 nt. A standard curve with serial dilutions of the linearized plasmid ranging from 4 x 10^2^ to 4 x 10^7^ copies per reaction was used to determine the absolute number of cDNA transcripts.

### Immunohistochemistry, imaging, and analysis

40μm coronal brain sections were cut using a cryostat (Leica Microsystems, Wetzlar, Germany) and collected into 96 well-plates containing cryoprotectant (30% ethylene glycol, 15% sucrose in TBS). Staining of free-floating sections was done in a 96 well plate, and samples were randomized and coded to create blinded conditions for analysis. Sections were blocked in 5% normal goat serum (NGS), 2% bovine serum albumin (BSA), 1% glycine, and 0.1% Triton X-100 in PBS. The primary antibodies against LAMP1 (Rat 1:1000, BD San Diego, CA, Cat ID: 553792) and GFAP (Mouse, 1:1000; Cell Signaling Technology, Danvers, MA, Cat. ID: 3670S) were diluted in 5% NGS and 2% BSA and applied overnight at 4°C. The next day, sections were incubated in secondary antibodies goat-anti-rat AlexaFluor 633 (1:500; Invitrogen, Eugene, OR) or goat-anti-mouse AlexaFluor 555 (1:500; Invitrogen, Eugene, OR) in 1% NGS for 1-2 hours at room temperature, and mounted onto glass SuperFrost plus slides (Fisher Scientific, Pittsburgh, PA) with Immu-Mount (Fisher Scientific, Pittsburgh, PA). Images were acquired using DM8i Leica Inverted Epifluorescence Microscope with Adaptive Focus (Leica Microsystems, Buffalo Grove, IL), Hamamatsu Flash 4.0 camera and advanced acquisition software package MetaMorph 4.2 (Molecular Devices, LLC, San Jose, CA) with an automated stitching function. The exposure time was kept the same for all sections within the same IHC experiment. Image analysis was performed using Fiji software (NIH, Bethesda, Maryland). Particle analysis and percentage-of-area measurements for LAMP1 and GFAP images were made after the same thresholding settings were applied to all images. The image analysis was done by an investigator blinded to the genotype and treatment groups. All images were decoded after the measurements were taken. Area and particle size values were averaged per genotype/treatment group and compared between groups using ordinary one-way ANOVA test and Dunnett’s correction for multiple comparisons in the GraphPad Prizm v9 software.

### Mass spectrometry proteomic analysis

Protein extraction from cerebral cortex, TMT labeling and LC-MS/MS analysis was performed as previously described (29). Raw data were submitted for analysis in Proteome Discoverer 3.0.1.23 (Thermo Scientific) software with Chimerys (MSAID, Germany). Assignment of MS/MS spectra was performed using the Sequest HT algorithm and Chimerys by searching the data against a protein sequence database including all entries from the Mouse Uniprot database (SwissProt 19,768 2019) and other known contaminants such as human keratins and common lab contaminants. Sequest HT searches were performed using a 20 ppm precursor ion tolerance and requiring each peptides N-/C termini to adhere with Trypsin protease specificity, while allowing up to two missed cleavages. 18-plex TMT tags on peptide N termini and lysine residues (+304.207146 Da) was set as static modifications and Carbamidomethyl on cysteine amino acids (+57.021464 Da) while methionine oxidation (+15.99492 Da) was set as variable modification. A MS2 spectra assignment false discovery rate (FDR) of 1% on protein level was achieved by applying the target-decoy database search. Filtering was performed using a Percolator (64bit version, reference 1). For quantification, a 0.02 m/z window centered on the theoretical m/z value of each of the six reporter ions and the intensity of the signal closest to the theoretical m/z value was recorded. Reporter ion intensities were exported in result file of Proteome Discoverer 3.0 search engine as an excel table. The total signal intensity across all peptides quantified was summed for each TMT channel, and all intensity values were adjusted to account for potentially uneven TMT labeling and/or sample handling variance for each labeled channel.

PSM-level data from each TMT channel was analyzed in R. The initial preprocessing involved missing value imputation using random forest and the removal of PSMs with incomplete or multiple protein mapping or isolation interference >70. The preprocessed data was aggregated to the protein level and normalized via Variance Stabilizing Normalization. Differential expression analysis was performed using the R package limma to compare proteomic profiles of untreated *Mcoln1^-/-^* cortical samples to wild-type and AAV-MCOLN1 treated *Mcoln1^-/-^* mice. Visualization was done through volcano plots, highlighting proteins with a significant difference (p < 0.05) and a log fold-change > |0.5|. Significantly altered proteins in the *Mcoln1^-/-^* vs. wild-type comparison were utilized to create a heatmap showing abundance differences across all samples in three experimental groups.

## Acknowledgments

Funding for this work was provided via the MGB Innovation Discovery Grant Award to Y.G and F.B. The authors are grateful to Drs. Sue Slaugenhaupt, Albert Misko and Patricia Musolino for fruitful discussions of this work and to the MLIV foundation, led by Randy Gold and Dr. Rebecca Oberman, for the tireless efforts supporting preclinical and clinical research of mucolipidosis IV.

## Author Contributions

M.S.- Investigation, Methodology, Formal Analysis, Writing – original draft, Writing – review & editing; M.B.- Investigation, Formal Analysis; Y.Y.- Investigation, Methodology; J.F.- Investigation, Methodology; S.S.- Formal Analysis, Data curation, Visualization; M.M. - Investigation, Formal Analysis; A.C. - Investigation, Formal Analysis, Writing – review & editing; B.B.- Investigation, Formal Analysis, Methodology; F.B. - Funding acquisition, Resources, Writing – review & editing; Y.G. – Conceptualization, Formal Analysis, Supervision, Visualization, Writing – original draft, Writing – review & editing.

## eTOC Synopsis

Mucolipidosis IV (MLIV) is a rare neurologic disease of childhood with extremely high unmet need. Here Grishchuk, Bei and colleagues show that AAV-CPP16-mediated gene transfer of the MLIV causative *MCOLN1* gene to symptomatic MLIV mice restores neurofunction, delays paralysis and alleviates brain pathology, indicating a promising therapeutic approach for clinical development.

