## supplemental info and figures for "A blood-brain-barrier penetrant AAV gene therapy rescues neurological deficits in mucolipidosis IV mice"

### Supplementary Information

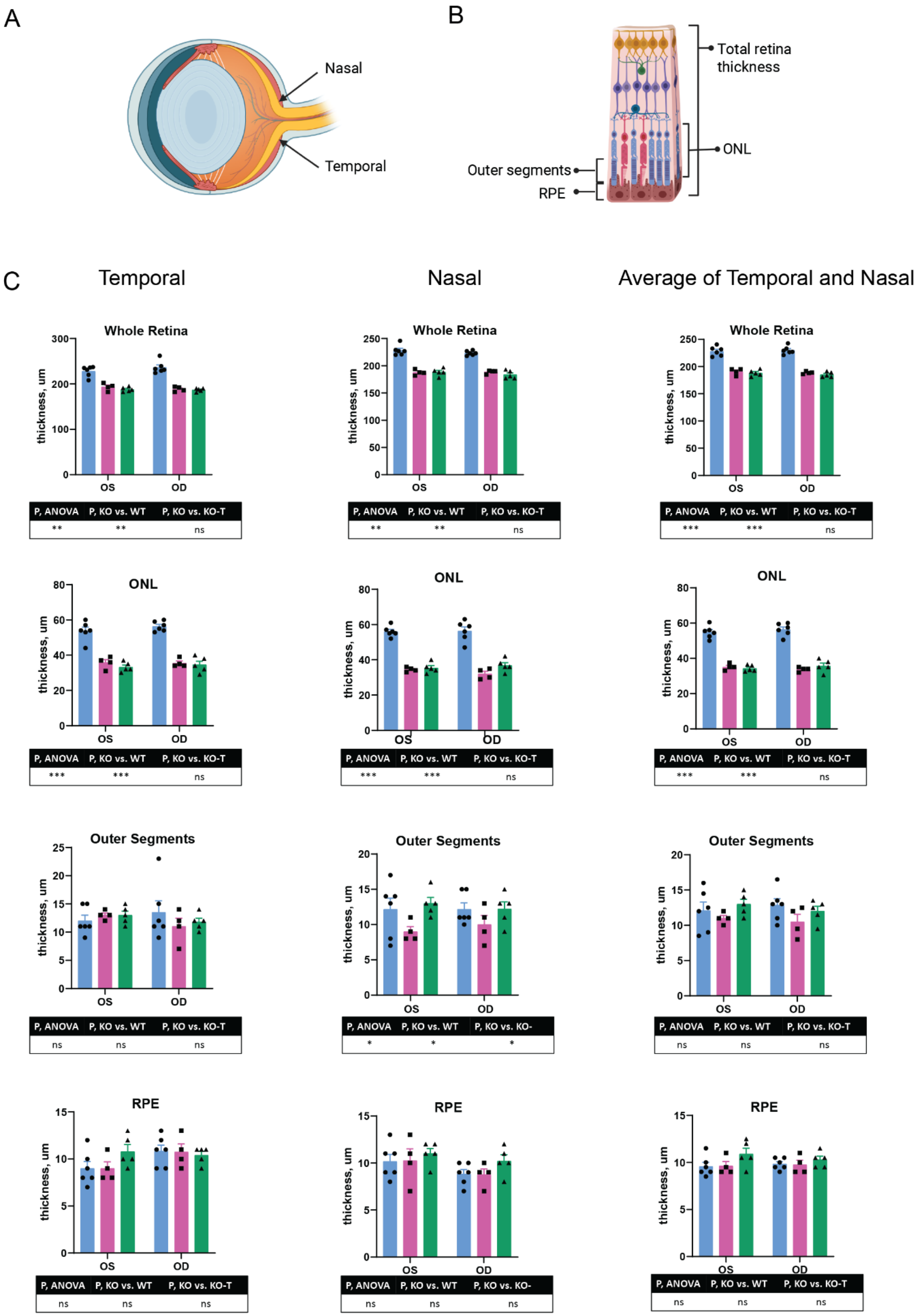
A
Nasal
Temporal
B
Total retina thickness
ONL
Outer segments
RPE
C
Temporal
Nasal
Average of Temporal and Nasal
Whole Retina
thickness, um
OS
OD
P, ANOVA | P, KO vs. WT | P, KO vs. KO-T
** | ** | ns
** | ** | ns
*** | *** | ns
ONL
P, ANOVA | P, KO vs. WT | P, KO vs. KO-T
*** | *** | ns
*** | *** | ns
*** | *** | ns
Outer Segments
P, ANOVA | P, KO vs. WT | P, KO vs. KO-T
ns | ns | ns
P, ANOVA | P, KO vs. WT | P, KO vs. KO-
* | * | *
P, ANOVA | P, KO vs. WT | P, KO vs. KO-T
ns | ns | ns
RPE
P, ANOVA | P, KO vs. WT | P, KO vs. KO-T
ns | ns | ns
P, ANOVA | P, KO vs. WT | P, KO vs. KO-
ns | ns | ns
P, ANOVA | P, KO vs. WT | P, KO vs. KO-T
ns | ns | ns

**Figure S1.**

**Figure S1. Systemic administration of CPP16-*MCOLN1* in young adult symptomatic *Mcoln1*$^{-/-}$ mice did not improve retinal thickness. A.** Schematic representation of the mouse eye, showing positions "nasal" and "temporal" where retinal measurements took place. **B.** Schematic presentation of the retinal structure showing acquired retinal measurements. **C.** Retinal layer thickness in WT- saline (blue), *Mcoln1*$^{-/-}$ - saline (pink) and *Mcoln1*$^{-/-}$- CPP16-*MCOLN1* (green) temporal and nasal sections of retina in the left (OS) and right (OD) eyes. Individual, group mean, and SEM values are presented, n=6 (WT saline), n=4 (KO saline), n=5 (KO CPP16-*MCOLN1*). Statistical analysis was done using one-way ordinary ANOVA and multiple comparison test using GraphPad Prizm v.9.

**Figure S2.**

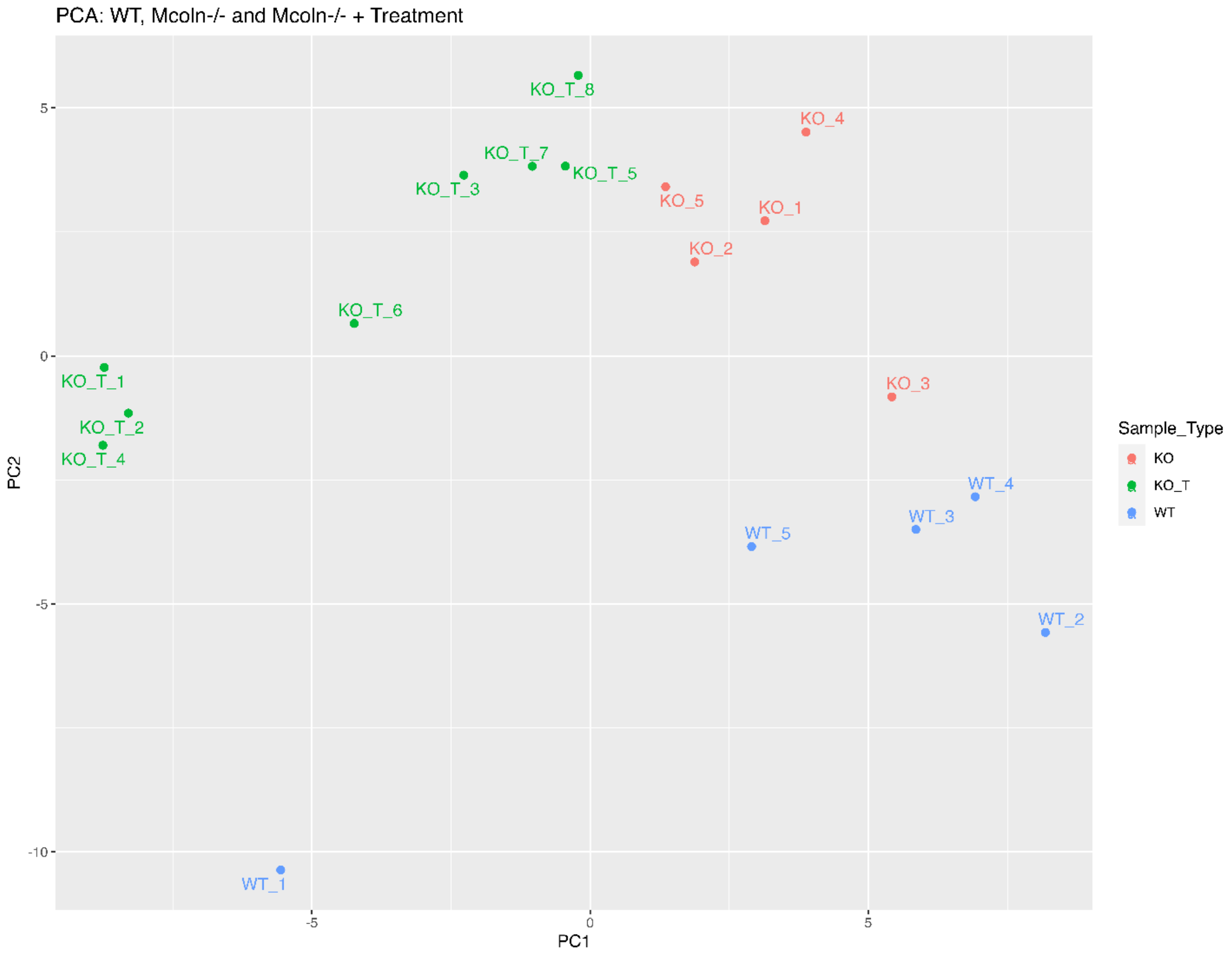

**Figure S2.** Principal component analysis (PCA) showing separation of *Mcoln1*$^{-/-}$ -saline (KO), *Mcoln1*$^{-/-}$ -CPP16-*MCOLN1* (KO-T) and WT-saline (WT) whole cortical homogenate samples via LC-MS/MS.

**Figure S3.**

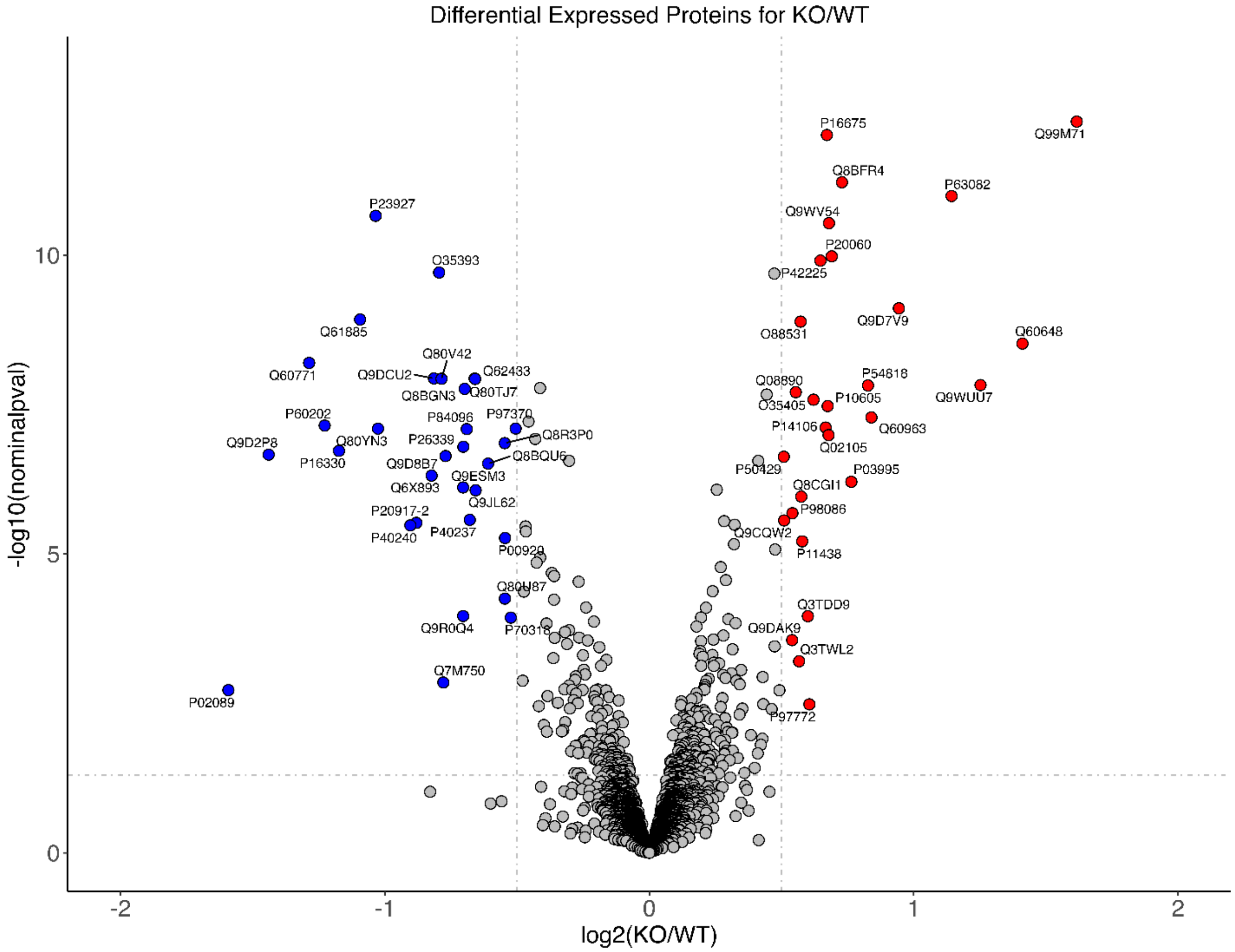


**Figure S3. Volcano plot showing upregulated and downregulated proteins in whole cortical homogenates from *Mcoln1*$^{-/-}$ and WT mice; log10 p<1.3 (p<0.05), log2FC>0.5.**

**Table S1. Protein abundances in whole cortical homogenates in *Mcoln1*$^{-/-}$ -saline, *Mcoln1*$^{-/-}$ -AAV-CPP16-*MCOLN1* and WT-saline mice.**

**Table S2. Common contaminants excluded from the analysis.**

**Table S3. UP and DOWN regulated proteins in whole cerebral cortex homogenates from WT and *Mcoln1*$^{-/-}$ saline-treated mice.**

**Table S4. UP and DOWN regulated proteins in whole cerebral cortex homogenates from *Mcoln1*$^{-/-}$ -CPP16-*MCOLN1* and *Mcoln1*$^{-/-}$ -saline mice.**
